# Sample-specific protein-protein interaction networks inferred from transcriptomics and proteomics show high similarities

**DOI:** 10.64898/2026.08.09.743737

**Authors:** Enikő Zakar-Polyák, Csaba Kerepesi

## Abstract

Contextualized protein-protein interaction networks provide crucial insight into diseases and other biological processes, but for a profound understanding of such processes and their distinct effects on individuals, the protein-protein interactions within individual samples must be investigated. A straightforward approach to estimate the PPI network of a sample is to restrict a general network of known PPIs to the proteins that are found in the sample. Although proteomics methods are becoming more accessible and precise, large-scale and single-cell studies still mainly target characterizing the transcriptomics profile of the samples, which is then often used as an approximation of the protein activities. The correlation of gene expression and protein abundance has been addressed in the past, but information about the deviations of the different omics-based estimates of the PPI networks is still lacking. In this study, we performed a comparative analysis of transcriptomic-based and proteomic-based sample-specific PPI network estimates to fill this gap. We created a framework for a comprehensive and transparent comparison of the two omics levels in two independent datasets, with a special focus on time-related network dynamics. We found that the size-adjusted characteristics of the different omics-based networks are very similar; the overall trend of how they change with time is also often the same, but the rate of the changes typically differs. The characteristics of the nodes present in both types of networks also show high similarity and often different time-related rates of change, but this varies among metrics. These results shed light on the properties of PPI network estimations and advise caution in interpreting them appropriately.

## 1 Introduction

Proteins are among the most essential building blocks of living organisms, establishing and controlling cellular processes through interactions. To have an adequate understanding of the inner biological activities of any organism, a systems-level view of the protein-protein interactions (PPIs) is necessary (Barabási & Oltvai 2004). Numerous methods, such as the yeast two-hybrid technology (Fields and Song 1989; Walhout and Vidal 2001), affinity purification mass spectrometry (Gavin et al. 2002; Gingras et al. 2007), and Co-Immunoprecipitation (Kessler 1975; Selbach and Mann 2006) have been developed to date to discover PPIs, accelerating the generation of high-quality experimental data. Then, in recent years, a great deal of effort was put into creating extensive databases of already known and presumed PPIs, providing a comprehensive map of the PPI network (Szklarczyk et al. 2023; Oughtred et al. 2021; Fischer et al. 2025; Veres et al. 2015).

Available protein-protein interaction networks mostly provide general information about interactions, which must be contextualized to yield meaningful insights into diseases and other biological processes. Restricting the network to the genes differentially expressed between conditions and examining the characteristics of this subnetwork and its nodes (i.e., proteins) has been performed to get a better understanding of diseases (Karimizadeh et al. 2019; Lee et al. 2011) or some aspects of aging (Wan et al. 2023; Altab et al. 2025). Instead of gene expression, protein abundance data can also be used to find the proteins exhibiting differences through processes such as aging, and explore their connections in the general PPI (Jin et al. 2025). Another study has created soybean tissue-specific PPI networks using multiple methods of combining transcriptome data and a general PPI network (J. Wang et al. 2019). Even ready-to-use tools have been created, which make the generation of context-specific PPI networks more accessible (Rosenberger et al. 2020; Do et al. 2024).

Although disease-specific, aging-specific, and tissue-specific networks provide valuable knowledge, more detailed information can be accessed through sample-specific networks. Some studies have created a network for each cancer sample by defining a set of genes of interest by comparing the cancer sample to a control group, and mapping the genes to a general PPI network (Liu et al. 2019; Zhang et al. 2022). Another way of creating sample-specific networks is to restrict a general network, e.g., a PPI network, to the genes expressed in a sample, resulting in a subnetwork of the general one, representing the sample. Another study created not sample-specific but age group-specific networks by integrating all samples belonging to a given age and defining a gene as being expressed at that age using the majority rule (Faisal and Milenković 2014). Most attempts to generate sample-specific PPI networks rely on transcriptomic data due to its comprehensive nature and accessibility; however, it is unclear how well gene expression predicts protein abundance (Laman Trip et al. 2025). To overcome the limitations of gene expression, proteomics data can be coupled with a PPI network instead (C. Wang et al. 2023), albeit with a potential loss of coverage. Another work produced a reference network of protein-protein associations, instead of interactions, also utilizing proteome data (Laman Trip et al. 2025).

Restricting the general PPI network to genes or proteins found in a sample, using transcriptomic or proteomic data, gives two different approximations of the true in vivo interaction network of the sample. The gene expression-based PPI network can be considered as an upper estimate of the true PPI network – in case of a complete general network of all possible PPIs – since, while transcriptomic measurements have high coverage, transcript levels are considered insufficient predictors of proteins, often over-predicting protein abundance (Choquet et al. 2025). The protein abundance-based PPI network, on the other hand, provides a lower estimate of the true in vivo interaction network, because the current proteomics experiments often still have technical or cost-related limitations on coverage (Messner et al. 2023). Then we ask the question of how similar the two different approximations are, with a special focus on time-related dynamics. We hypothesize that their properties exhibit similar time-related changes; therefore, both estimates can be utilized to reveal time-related (e.g., age) dynamics of the PPI in samples.

In this work, we explore this question by performing a comparative analysis of sample-specific transcriptomic-based and proteomic-based protein-protein interaction networks. We generate networks using matched multi-omics measurements at different time points to minimize batch effects and utilize high-confidence, known physical PPIs from the STRING database (Szklarczyk et al. 2023). We mainly focus our analysis on time-related network dynamics as a useful application of sample-specific networks. This work aims to provide insight into how the choice of the omics level affects the dynamics of the inferred PPI networks, draw attention to the limitations of network estimations, and guide future work in making methodological choices appropriate for the purpose of the work.

## 2 Methods

### 2.1 Datasets

We used two multi-omics datasets containing both transcriptomic and proteomic measurements from the same samples to create the sample-specific networks. From the dataset of Lu et al., we used the data of the human hepatocellular cell line MHCC97H (Lu et al. 2023). Gene expressions were measured at the first 9 culturing generation phases; matched proteomic measurements were taken at 8 generation points, excluding the second generation, compared to the transcriptomic measurements. We used the available processed transcriptomic data containing the RPKM values of the genes and the processed output of the DDA mass spectrometry analysis. In total, 9 transcriptomic and 8 proteomic sample-specific networks were generated, and for the direct comparison of the matched samples, the second-generation transcriptomic data were excluded.

The dataset presented by Becker et al. and Casas-Vila et al. contains paired transcriptomic and proteomic data measured at 14 time points during Drosophila melanogaster embryogenesis (Becker et al. 2018; Casas-Vila et al. 2017). At each time point, four replicates were measured in the samples, resulting in 56 transcriptomic and 56 proteomic measurements in total. We used the available processed count per gene files and the processed *proteinGroups* file of the DDA mass spectrometry analysis.

To construct our sample-specific networks, we utilized the STRING v12.0 database as general PPI networks (Szklarczyk et al. 2023). We used the Homo sapiens physical subnetwork for the cell line dataset, and the Drosophila melanogaster physical subnetwork for the embryogenesis dataset (Fig. 1a).

**Figure 1.**
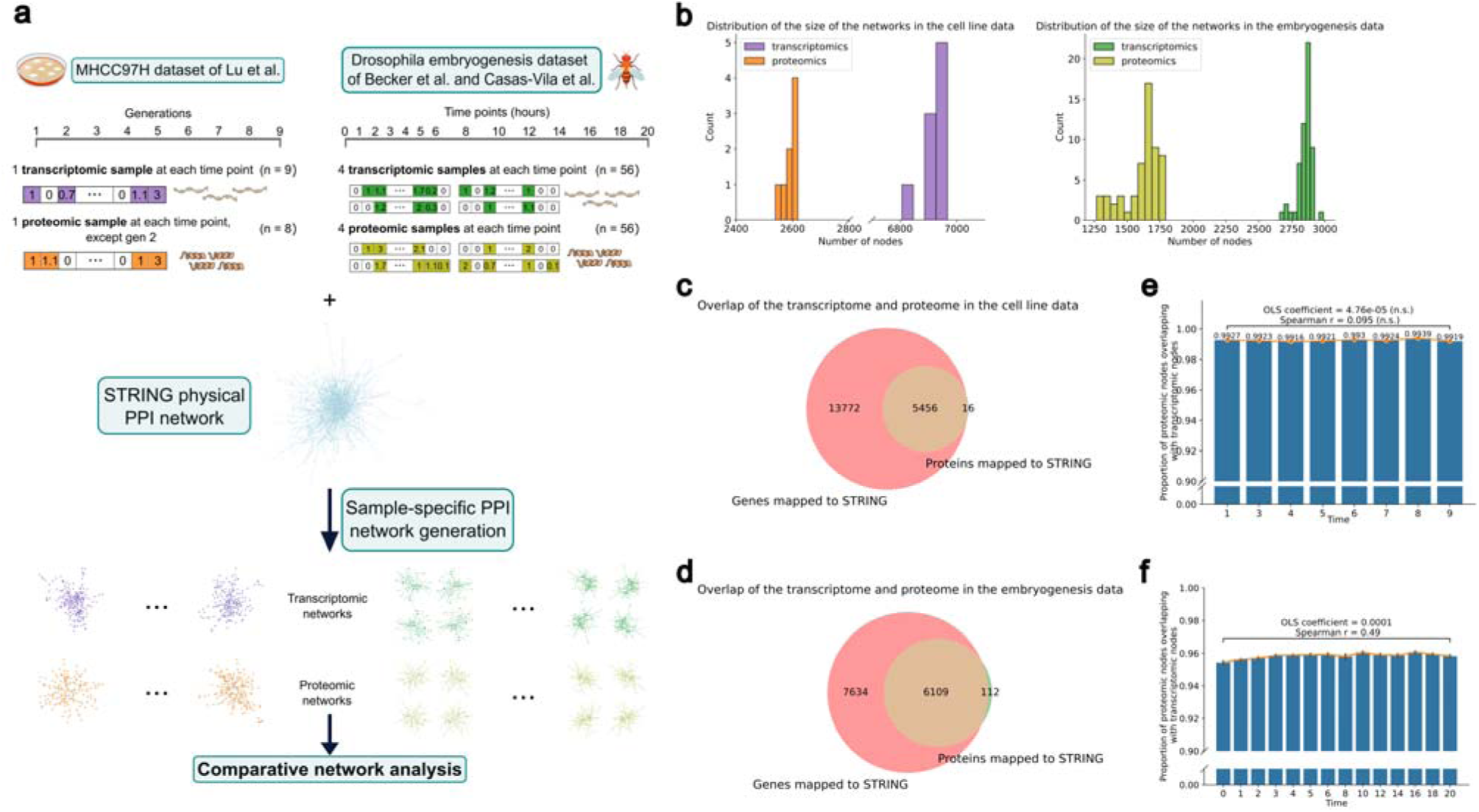
Schematics of the study and overlap analysis of the networks inferred from transcriptomic or proteomic data. (a) Schematics of the study. (b) Comparison of the number of nodes in the networks. (c) Overlap (brown) of the genes in the transcriptomic data mapped to STRING (red) and the proteins in the proteomic data mapped to STRING (green) in the cell line dataset. (d) Overlap (brown) of the genes in the transcriptomic data mapped to STRING (red) and the proteins in the proteomic data mapped to STRING (green) in the embryogenesis dataset. (e) The proportion of nodes in the proteomic-based networks overlapping with the nodes in the transcriptomic- based networks for each time point in the cell line dataset. An OLS model was fitted to the proportion value as a function of time, and the coefficient of the time variable in the model is shown in the plot, as well as the Spearman correlation coefficient between the time and the proportion value. (f) The proportion of nodes in the proteomic-based networks overlapping with the nodes in the transcriptomic-based networks for each time point in the cell line dataset. The variation of the samples at each time point is also shown as an error bar. The coefficient of time in the OLS model fitted to the proportion value as a function of time is shown in the plot as well as the Spearman correlation coefficient between time and the proportion value. The model coefficient and the correlation coefficient significantly differed from zero. Significance was determined at p<0.05.

### 2.2 Generation of sample-specific PPI networks

To map the sample-specific data to the general PPI network, the gene names or identifiers in the transcriptomic datasets were matched to the STRING protein identifiers through the preferred name column of the protein info file available in the STRING database. Non-matched genes were further checked and mapped through the protein aliases file, also available in the database. Then, the general physical network was restricted to the genes with a non-zero RPKM value in the given sample. Note that we chose zero as the threshold not to exclude genes with low expression either. Only the high- confidence connections, with a score larger than 0.700 for the experiments or database source, were retained in the network.

From the proteomic datasets, the reverse and contaminated entries were excluded, as well as protein groups having more than one majority protein. Since in the Drosophila dataset, transcripts are mapped to protein groups, the data entries with multiple transcripts mapping to the same protein were retained. Then, the protein names or identifiers were mapped to the STRING identifiers using the preferred names in the protein info file and the aliases file, similarly to before. One-to-many assignments were manually checked and corrected based on the UniProt and GeneCards databases. Sample-specific PPI networks were generated by restricting the general network to the proteins with non-zero LFQ intensity values in the given sample. In the Drosophila dataset, if more than one transcript group mapped to the same protein, the restriction was based on the maximum value of these groups. Similarly as before, only the high-confidence connections were retained in the network.

Note that we constructed a network for each available datapoint in the datasets, meaning that in the embryogenesis dataset, separate networks were created for each replicate too.

### 2.3 Characterization of networks

To compare the different omics-based sample-specific networks, we first aimed to thoroughly characterize them. We collected and calculated an extensive set of network metrics capturing various properties, including size-, density-, and centrality-related measurements, as well as network entropy metrics (Dong and Horvath 2007; Sun and Wandelt 2014; Zakar-Polyák et al. 2023). The complete set of metrics can be found in Table 1.

**Table 1.** Network metrics calculated in the study and the corresponding scaling constants.

| Notation | Description | Scaling constant |
| --- | --- | --- |
| num_nodes | Number of nodes, denoted by $N$ | |
| num_edges | Number of edges | $\frac{N \times (N - 1)}{2}$ |
| num_components | Number of components | $N - 1$ |
| avg_deg | Average degree | $N - 1$ |
| max_deg | Maximum degree | $N - 1$ |
| density | <sup>1</sup>Density defined as $\frac{\text{avg\_deg}}{N-1}$ | 1 |
| centralization | <sup>1</sup>Centralization defined as $\left(\frac{\text{max\_deg}}{N-1} - \text{density}\right) \times \frac{N}{N-2}$ | 1 |
| heterogeneity | <sup>1</sup>Heterogeneity defined as $\frac{\sqrt{\text{variance}(\text{degree})}}{\text{avg\_deg}}$ | $\frac{\sqrt{2}}{2} \times \sqrt{N-2}$ |
| glob_clust | Global clustering coefficient defined as $3 \frac{\text{number of triangles}}{\text{number of triads}}$ | 1 |
| avg_clust | Average clustering coefficient* | 1 |
| max_clust | Maximum clustering coefficient* | 1 |
| skew_deg_dist | Skewness of the degree distribution | $\frac{2(N-2)}{\sqrt{N-1}}$ |
| assortativity_degree | <sup>2</sup>Assortativity coefficient | 2 |
| entropy_shannon | Shannon entropy of the degree distribution | $-\log_2 \frac{1}{N}$ |
| entropy_rw | <sup>3</sup>Random walker entropy defined as $\frac{1}{N \ln(N-1)} \sum_{i=1}^N \ln \text{degree}_i$ | 1 |
| entropy_ks | <sup>4</sup>Kolmogorov-Sinai entropy defined as the logarithm of the maximum eigenvalue of the adjacency matrix | $\ln(n-1) - 1$ |
| node_bc_max | Maximum node betweenness centrality** | 1 |
| node_bc_avg | Average node betweenness centrality** | 1 |
| edge_bc_max | Maximum edge betweenness centrality*** | 1 |
| edge_bc_avg | Average edge betweenness centrality*** | 1 |

<sup>1</sup>(Dong and Horvath 2007) <sup>2</sup>(Newman 2002) <sup>3</sup>(Small 2013) <sup>4</sup>(Demetrius and Manke 2005)
\*Clustering coefficient of node $u$ is defined as $\frac{2 \times \text{number of triangles through node } u}{\text{degree}_u \times (\text{degree}_u - 1)}$ .
\*\*Node betweenness centrality of node $u$ is defined as $\frac{\text{number of shortest paths through node } u}{\text{number of shortest paths}}$ .
\*\*\*Edge betweenness centrality of edge $e$ is defined as $\frac{\text{number of shortest paths through edge } e}{\text{number of shortest paths}}$ .

Addressing the inherent differences in the metric values caused by the size discrepancy of the networks is essential for a proper comparison of the network dynamics (Fig. 1b). To make the metrics size-independent, we scaled them by dividing by the theoretical range of the metric, given the number of nodes in the network. For the skewness of the degree distribution, which has no theoretical minimum or maximum value, we utilized proven lower and upper bounds to define the range (Wilkins 1944; Kirby 1974). The scaling constants used can also be found in Table 1. This way of rescaling also ensures that the difference between the two networks’ metric values is always between 0 and 1.

### 2.4 Comparison of time-related dynamics

To assess the similarity between the time-related dynamics of the transcriptomic-based and proteomic-based networks, we performed two complementary analyses. First, we fit a unique linear regression model to the metric value with time as the independent variable for the transcriptomic-based and the proteomic-based data, and compare the slope of the two models. If both slopes significantly differ from zero and have the same direction, i.e., both are positive or negative, or neither of them differ significantly from zero, then we conclude that the given metric follows the same time-related trend in both data.

For the second analysis, we fit a linear regression model to the metric value on the transcriptomic-based and proteomic-based data together with the time and the source (transcriptomic or proteomic) as the independent variables and an interaction term between them. If the interaction term does not differ significantly from zero, meaning that parallel regression lines can be fitted to the two data separately, then we conclude that the given metric has the same time-related rate of change in both data. We used the Freedman-Lane permutation method to determine whether the interaction term significantly differs from zero, because it provides a robust, non-parametric framework to test a variable in a multivariate linear regression model (Freedman and Lane 1983; Winkler et al. 2014).

### 2.5 Comparison of node-level characteristics

Besides the whole network-level characterization of the sample-specific networks, we also calculated node-level metrics to capture further details of the networks. Given the network-level metrics, we calculated the degree, the clustering coefficient, and the node betweenness centrality for each node in each network. To make the metrics independent of the number of nodes, we scaled them according to Table 1, namely, divided the degree by N-1. Note that the scaled degree equals to the degree centrality measurement. To further compare the different omics-based sample-specific networks based on the node-level properties, we selected the common nodes, i.e., nodes in at least one transcriptomic-based and at least one proteomic-based network in the given dataset. This yielded 3019 nodes in the cell line dataset and 2079 nodes in the embryogenesis dataset. We compared the time-related dynamics of the characteristics of each common node as described in Section 2.4.

### 2.6 Statistical analysis and reproducibility

Significance was determined at p<0.05 in the cell line dataset due to the small sample size and at p<0.05 after Bonferroni correction for the number of metrics in the embryogenesis dataset in the network-level analyses. Significance was determined at p<0.05 after FDR correction in both datasets in the node-level analyses.

All analyses were performed using Python 3.8.17. The essential packages with the version numbers in parentheses are listed below:

- Packages for data manipulation: numpy (1.24.4), pandas (1.5.3)
- Package for network analysis: networkx (3.1)
- Statistics packages: scipy (1.10.1), statsmodels (0.14.0)
- Packages for figure generation: matplotlib (3.3.4), seaborn (0.11.2).

The data generated in this work is available on Zenodo at <u>10.5281/zenodo.21129948</u>. The code generated in this work is available on GitHub at https://github.com/polyake/sample_specific_ppi.

## 3 Results

We performed a comparative analysis of sample-specific PPI networks generated from transcriptomic or proteomic measurements to assess the similarity of the two omics in terms of the dynamics of the inferred PPI networks (Fig. 1a). We generated sample-specific networks by taking the subnetwork of the general PPI network induced by the expressed genes and the abundant proteins in the sample and calculated an extensive set of metrics to characterize the structure of the networks. We performed a direct comparison of the metric values and compared the time-related trajectories as well.

### 3.1 Large overlap between the transcriptomic-based and the proteomic-based networks

To create a proper basis for our study, we explored the overlap between the transcriptomic and the proteomic data, as well as the nodes of the inferred networks. The inherent coverage difference between the transcriptomic and proteomic measurements is directly visible in the dimensions of the different omics data. In the cell line dataset, 28 252 genes are present in the measured transcriptome, and 19 228 of these genes map to the STRING PPI network, while there are 5 510 protein measurements in the proteome after quality control, and 5 472 unique proteins map to the STRING PPI network. In the embryogenesis dataset, 17 558 genes are measured in the transcriptome, and 13 743 genes map to the STRING PPI network, while 6 915 measurements are present in the proteome after quality control and 6 221 unique proteins map to the STRING PPI network. Due to the inherent coverage difference, the difference in the size (i.e., the number of nodes) of the sample-specific networks inferred from the different omics data is also notable (Fig. 1b). In the cell line dataset, the number of nodes in the transcriptomic-based networks ranges from 6 806 to 6 967 and from 2 539 to 2 619 in the proteomic-based networks. The size of the transcriptomic-based networks in the embryogenesis dataset ranges from 2 657 to 2 989 and from 1 268 to 1 792 in the proteomic-based networks. The other raw, non-scaled network metric values also often show large differences – although they are often of similar magnitude - between the transcriptomic-based and proteomic-based networks (Supplementary Fig. 1a and b), while the metrics which are scaled by definition (e.g. betweenness centrality) are of the same magnitude. The node-level metric distributions, i.e. the distribution of metrics calculated for each node, largely overlap (Supplementary Fig. 1c and d), which further validates that the correction of the metrics is necessary if we aim to comparecharacteristics not affected by size differences.

While the coverage difference is remarkable, there is a large overlap between the mapped genes and proteins. In the cell line dataset, 99.7% of the mapped proteins measured in the proteome have a corresponding mapped gene in the measured transcriptome (Fig. 1c), and the overlap is 98.2% in the embryogenesis dataset (Fig. 1d). This suggests that the captured fragment of the proteome in the inferred PPI networks is roughly the subset of the measured set of genes in the respective inferred networks.

The overlap between the nodes of the transcriptomic-based and the proteomic-based networks is similarly large throughout the time points as well. In the cell line dataset, the proportion of the nodes in the proteomic-based networks overlapping with the nodes in the transcriptomic-based networks does not change significantly with time, it is constantly above 0.99 (Fig. 1e). While there is a significant, moderate relation between the time and the overlap in the embryogenesis dataset (Spearman r = 0.49), the change is subtle and the proportion of proteomic nodes present in the transcriptomic-based networks is always above 0.95 (Fig. 1f). The latter observation suggests that the composition of the transcriptomic-based and the proteomic-based PPI networks becomes more similar over time.

### 3.2 Weak correlation, but small absolute difference of metric values of transcriptomic-based vs proteomic-based PPIs

First, we assessed the relationship between the network characteristics of the transcriptomic-based and the proteomic-based networks as a preliminary comparison. We calculated the Spearman correlation coefficient between the two types of networks for each scaled metric. In the cell line dataset, the maximum clustering coefficient and the average node betweenness centrality show strong positive correlation, but the correlations are weak overall, even negative for 8 out of the 20 metrics (Fig. 2a). In the embryogenesis dataset, 7 out of the 20 metrics show strong correlation with a Spearman correlation coefficient above 0.6, however, negative correlations can also be observed for 8 metrics (Fig. 2b).

**Figure 2.**
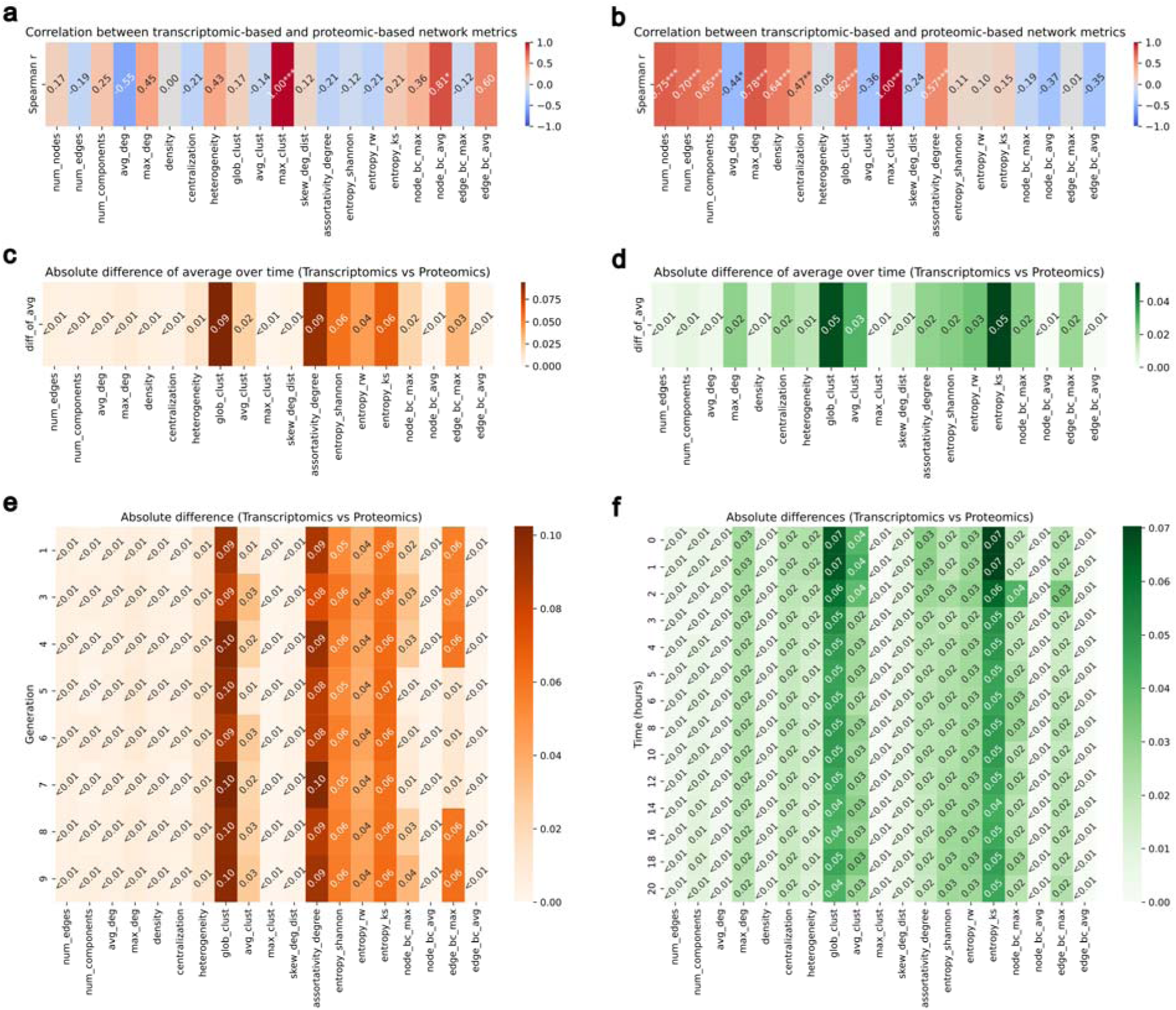
Comparison of the metric values of the networks inferred from transcriptomic and proteomic data. (a) Correlation between the scaled metric values of the transcriptomic-based and proteomic-based networks in the cell line dataset. Spearman correlation coefficient for each metric is shown in the heatmap. (b) Correlation between the scaled metric values of the transcriptomic-based and proteomic-based networks in the embryogenesis dataset. (c) Heatmap of the absolute differences between the average scaled metric values of the networks in the cell line dataset. The average was calculated over the whole set of transcriptomic- vs. proteomic-based networks. (d) Heatmap of the absolute differences between the average scaled metric values of the networks in the embryogenesis dataset. (e) Heatmap of the absolute differences between the scaled metric values of the transcriptomic-based vs. proteomic-based networks at each generation in the cell line dataset. (f) Heatmap of the absolute differences between the average scaled metric values of the transcriptomic- based vs. proteomic-based networks at each time point in the embryogenesis dataset.

**Figure 3.**
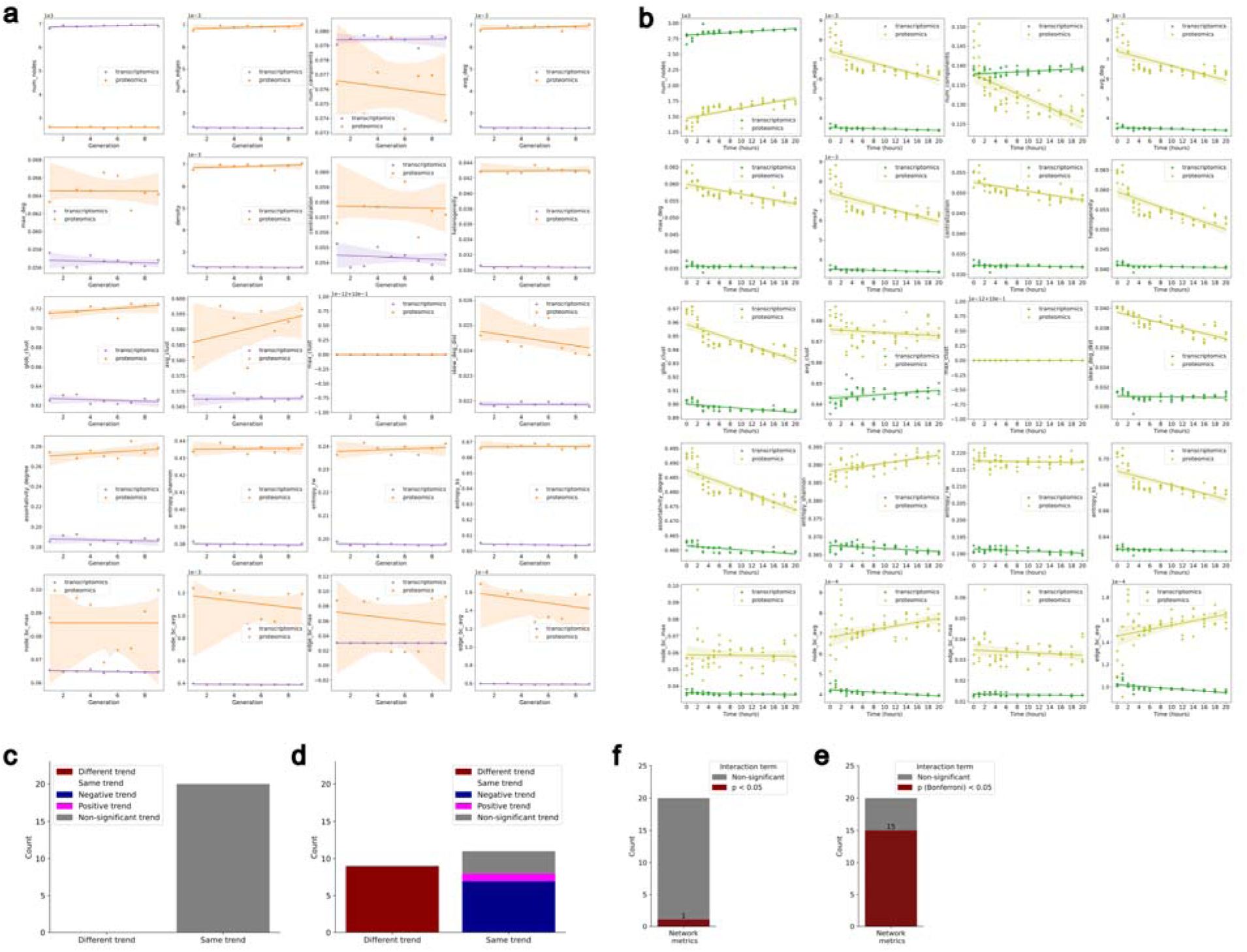
Comparison of the time-related changes of the characteristics of the networks inferred from transcriptomic and proteomic data. (a) The relation between time and scaled network metrics in the cell line dataset. The data points and the fitted linear regression lines are shown. n=9 for the transcriptomic data and n=8 for the proteomic data. (b) The relation between time and scaled network metrics in the embryogenesis dataset. n= 56 for the transcriptomic and n=56 for the proteomic data. (c) The comparison of the number of metrics with the same and different time-related trends in the transcriptomic-based and the proteomic-based networks in the cell line dataset. Separate linear regression models were fit to the scaled metric value for the transcriptomic-based and the proteomic-based data. If both slopes significantly differ from zero and have the same direction, then we conclude that the given metric follows the same time-related trend in both datasets. (d) The comparison of the number of metrics with the same and different time-related trends in the transcriptomic- based and the proteomic-based networks in the embryogenesis dataset. (e) The number of metrics with parallel time-related change in the transcriptomic-based and the proteomic-based networks in the cell line dataset. An OLS model with an interaction term was fitted to the scaled metric value as a function of time and source. Significance of the interaction term was determined by the Freedman- Lane permutation test at p<0.05 after Bonferroni correction. (f) The number of metrics with parallel time-related change in the transcriptomic-based and the proteomic-based networks in the embryogenesis dataset.

Then we performed another overall comparison of the network characteristics. We calculated the absolute difference between the average rescaled metric values of the transcriptomic-based and proteomic-based networks. The average was taken across the whole dataset for each metric, separately. A small difference between the averages can be observed in both datasets, with a maximum difference of 0.09 in the cell line dataset and 0.05 in the embryogenesis dataset (Fig. 2cd). In both datasets, the clustering-, assortativity-, entropy-, and betweenness centrality-related metrics show the highest, but still small differences.

We also compared the characteristics of the different omics-based networks at each time point. In this case, the average was taken across the networks corresponding to a given time point. We note that in the cell line dataset, this means direct comparison of the networks, because only one transcriptomic and one proteomic measurement is available at each time point. As before, the absolute differences are overall small in both datasets (maximum = 0.01), and there is no major time-related divergence of the metrics (Fig. 2ef). We would highlight the 5^th^-7^th^ generations in the cell line dataset, where the networks’ maximum node and edge betweenness centralities are noticeably more similar than in the other generations. In the embryogenesis dataset, the clustering coefficients and the Kolmogorov-Sinai entropy exhibit slight changes with time. Both characteristics of the networks become more similar after 3 hours of development.

Overall, while the network values do not correlate strongly between the two types of data, given that the difference between the rescaled values is interpreted in the 0-1 range, all the metrics showing differences smaller than 0.1 and of the same magnitude suggest that the PPI networks inferred from transcriptomic and proteomic data have very similar characteristics.

### 3.3 Different rates of change with time, but similar trends

We further compared the time-related dynamics of the networks inferred from transcriptomic and proteomic measurements by three complementary analyses.

To assess the time-related trend differences in the metrics, we fitted separate linear regression models to the characteristics of the transcriptomic-based and proteomic-based networks and compared the two slopes to determine whether both slopes significantly differ from zero and have the same direction, hence whether the metrics follow the same time-related trend in both datasets or not. In the cell line dataset, none of the metrics showed significant changes with time in either type of network, suggesting that the trend of the characteristics with time is similar in the transcriptomic-based and proteomic-based networks (Fig. 3ac, Supplementary Table 1). In the embryogenesis dataset, 11 out of the 20 metrics showed the same time-related trends in the different omics-based networks, meaning that either none of them changed significantly with time or both changed significantly in the same direction (Fig. 3bd, Supplementary Table 2). The metrics exhibiting the same trend include characteristics such as the number of nodes and edges, average degree and density, as well as the Kolmogorov-Sinai entropy, while the other entropy metrics and some betweenness centrality measurements have different time-related trends (Supplementary Table 2). Interestingly, 2 of the 20 metrics (Shannon-entropy and the number of components) show totally opposite trends (significant decrease vs significant increase).

For a more detailed comparison, we also analyzed the rate of change of the metrics with time by fitting a linear regression model to the metric on the whole data (both transcriptomic-based and proteomic-based network data) with an interaction term. In the cell line dataset, the interaction term was not significant for 19 of the 20 metrics, suggesting parallel slopes of the time-related changes of the different omics-based networks (Fig. 3af, Supplementary Table 1). In the embryogenesis dataset, however, the interaction term significantly differed from zero for 15 out of the 20 metrics, meaning that the rate of change cannot be considered the same (Fig. 3be, Supplementary Table 2). Only the average and maximum clustering coefficient, the random walker entropy, and the maximum node and edge betweenness centralities showed parallel changes with time.

Overall, the results suggest that while the rate of change of various network metrics with time often differs for the transcriptomic-based and proteomic-based networks, the trend of the changes is often still the same.

### 3.4 Varying similarity of different node characteristics

While whole-network metrics capture various characteristics and allow the comprehensive comparison of networks, important local structures and node-level discrepancies might stay hidden. Aiming to capture as much detail as possible, we also calculated node-level metrics and compared each node present in both transcriptomic-based and proteomic-based networks the same way as the whole networks.

First, we calculated the Spearman correlation coefficient between the transcriptomic-based and proteomic-based networks for each node and metric. In the cell line dataset, the distribution of the nodes’ correlation coefficients resembles a normal distribution for the clustering coefficient and the node betweenness centrality, meaning that most nodes’ metric values weakly correlate between the two types of networks with many negative correlation and a remarkable number of strong positive correlation (Fig. 4a). The distribution of the correlation coefficient for the nodes’ degree centrality has a notable peak at 0.2, but there are still many negative and positive correlations as well. In the embryogenesis dataset, most nodes’ degree centrality strongly correlates (Spearman r > 0.7), while the clustering coefficient weakly correlates (-0.25 < Spearman r < 0.25) between the two types, and the node betweenness centrality shows a mixed pattern (Fig. 4b).

**Figure 4.**
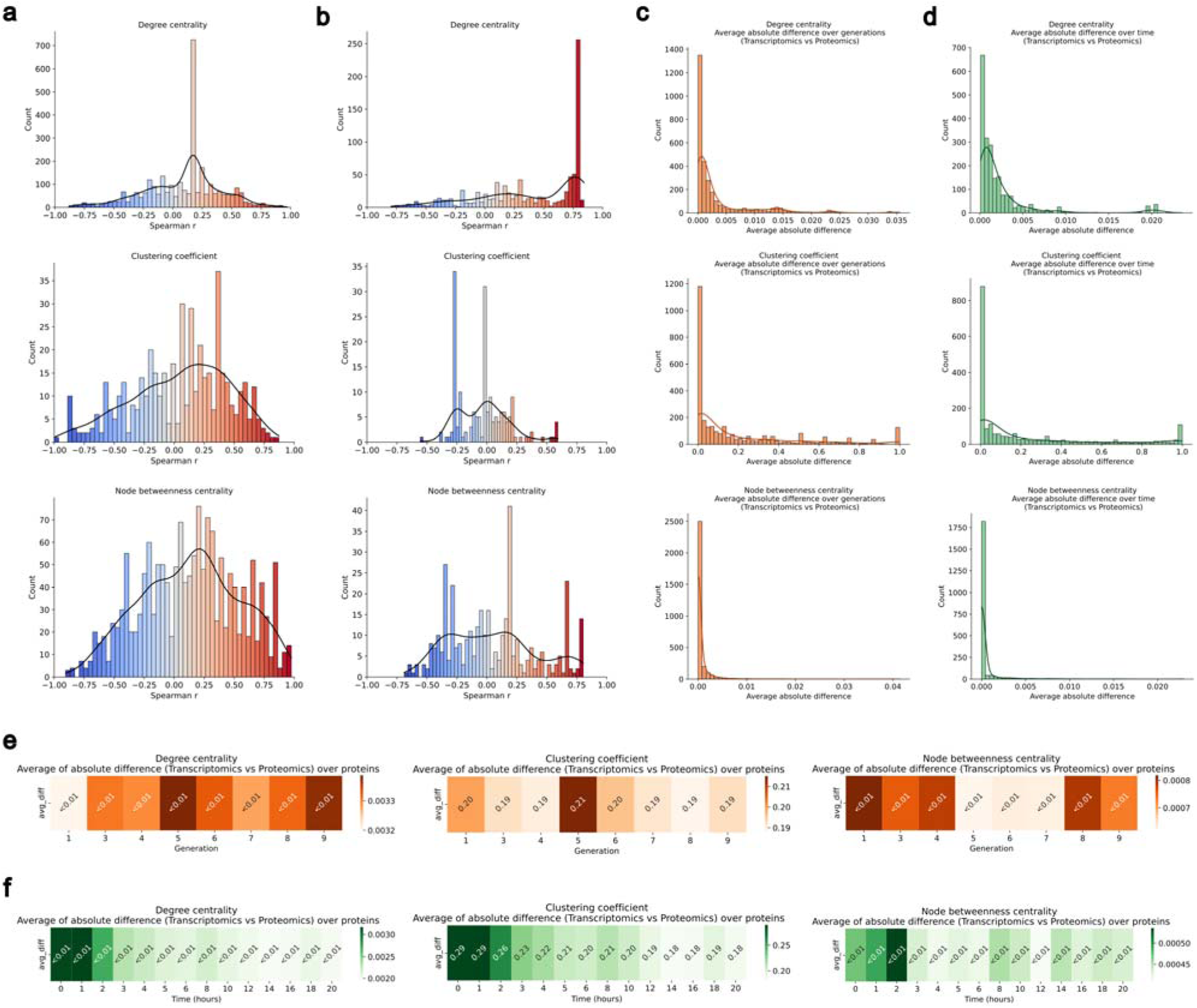
Comparison of the node-level metric values of the networks inferred from transcriptomic and proteomic data. (a) Distribution of the correlations between the node-level metric values of the transcriptomic-based and proteomic-based networks in the cell line dataset. Spearman correlation was calculated for each node common in the two types of networks. (b) Distribution of the correlations between the node-level metric values of the transcriptomic-based and proteomic-based networks in the embryogenesis dataset. (c) Distribution of the average absolute difference between the metric values of the transcriptomic-based and proteomic-based networks in the cell line dataset. Absolute difference was calculated for each node common in the two types of network, and the average was calculated over generations for each node. (d) Distribution of the average absolute difference between the metric values of the transcriptomic-based and proteomic-based networks in the embryogenesis dataset. Absolute difference was calculated for each node common in the two types of networks, and the average was calculated over time points for each node. (e) Heatmap of the average absolute differences between the metric values of the transcriptomic-based and the proteomic-based networks at each generation in the cell line dataset. Absolute difference was calculated for each node common in the two types of networks, and the average was calculated for each sample. (f) Heatmap of the average absolute differences between the metric values of the transcriptomic-based and the proteomic-based networks at each time point in the embryogenesis dataset.

We calculated the absolute difference of the rescaled node-level metrics for each common node between the transcriptomic-based and proteomic-based networks and compared the distribution of the average differences over time as well as the average difference over nodes at each time point. In both the cell line and the embryogenesis datasets, the scaled degree and the betweenness centrality of the nodes are overall very similar: the maximum average absolute difference is below 0.04 for both metrics (Fig. 4c-f). The clustering coefficient of the nodes shows larger dissimilarity between the transcriptomic-based and the proteomic-based networks on average (0.19-0.21 in the cell line dataset, 0.18-0.29 in the embryogenesis dataset), and it ranges from 0 to 1 across nodes too (Fig. 4c-f). Since the clustering coefficient is sensitive to small changes, e.g., the deletion or appearance of a single connection, it is suitable to highlight even subtle differences. However, the majority of the nodes still exhibit an average absolute difference close to zero, suggesting overall high similarity of node properties. We highlight that in the embryogenesis dataset, there is a remarkable decrease in the average absolute difference of the clustering coefficient with time, with a major drop around 3 hours (Fig. 4f). This suggests that the nodes in the transcriptomic-based and proteomics-based networks become more similar over development time, even based on the sensitive-to-change clustering coefficient, which we previously observed on the network level too.

We also compared the time-related dynamics of the node-level properties in the network inferred from transcriptomic and proteomic data, the same way as before. First, we fitted separate linear regression models to each characteristic of each common node in the transcriptomic-based and proteomic-based networks and compared their slopes to assess time-related trend differences. In the cell line dataset, we can observe similar time-related trends in both types of networks for all nodes, since none of the nodes exhibit significant change with time (Supplementary Data). In the embryogenesis dataset, 36% of the common nodes’ degree centrality, 53% of the nodes’ clustering coefficient, and 74% of the nodes’ betweenness centrality show similar time-related trends (Fig. 5ab, Supplementary Data). The similar degree trends mostly consist of a significant decrease with time, while the similar clustering and node betweenness trends consist of non-significant changes (Fig. 5b).

**Figure 5.**
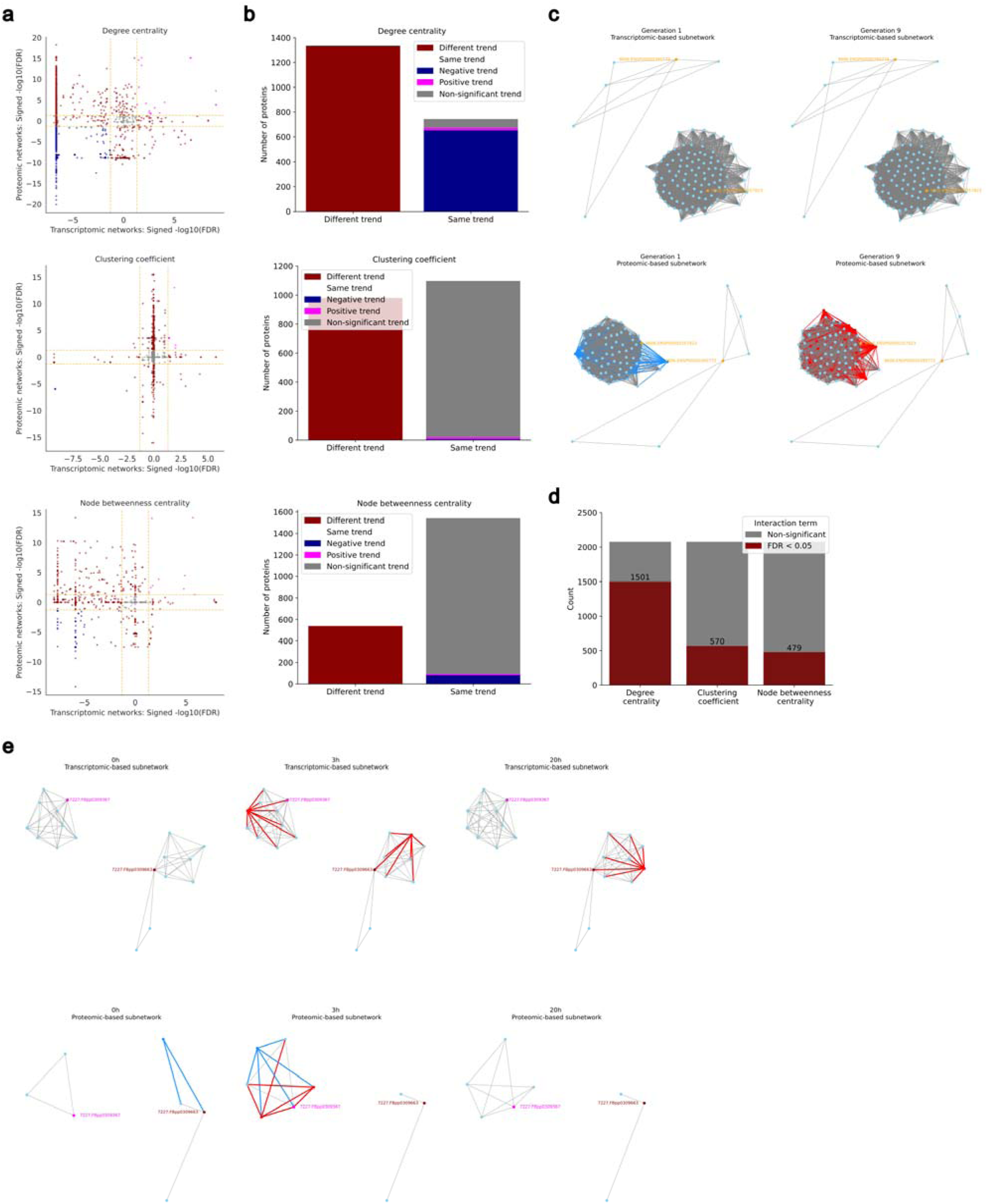
Comparison of the time-related changes of the node-level characteristics of the networks inferred from transcriptomic and proteomic data. (a) Comparison of the time-related trend of the node characteristics in the transcriptomic-based and proteomic-based networks in the embryogenesis dataset. Separate linear regression models were fitted to the metric values as a function of time on the transcriptomic-based and proteomic-based networks. The sign of the model coefficient was multiplied by -log(FDR), where FDR is the FDR-corrected p-value of the coefficient. This value for the transcriptomic-based networks was plotted against the value for the proteomic-based networks. Each point in the plot corresponds to a node in the networks. The horizontal and vertical yellow lines correspond to the significance threshold of FDR < 0.05. The points in the bottom left corner, the top right corner and the grey area in the middle have the same time-related trend in the two types of networks. (b) Summary of (a). The comparison of the number of nodes with the same and different time-related trends in the transcriptomic-based and the proteomic-based networks in the embryogenesis dataset. (c) Illustration of two subnetworks of a transcriptomic-based and a proteomic-based network at two time points in the cell line dataset. Two randomly selected nodes (orange) and their neighbors are shown. The edges and nodes disappearing in the later network are shown in blue, and the edges and nodes appearing compared to the earlier network are shown in red. (d) The number of nodes with parallel time-related change in the transcriptomic-based and the proteomic-based networks in the embryogenesis dataset. An OLS model with an interaction term was fitted to the metric value as a function of time and source on each common node. Significance of the interaction term was determined by the Freedman-Lane permutation test at FDR < 0.05. (e) Illustration of two subnetworks of a transcriptomic-based and a proteomic-based network at three time points in the embryogenesis dataset. The pink node (7227.FBpp0309367) is a protein with parallel time-related changes (increase) of its clustering coefficient in the transcriptomic-based and the proteomic-based networks. The dark red node (7227.FBpp0309663) is a protein with different time-related trends of its clustering coefficient in the two types of networks.

We assessed the difference between the rate of change of the node properties in the different omics-based networks by fitting a linear regression model to each metric of each node (including both transcriptomic-based and proteomic-based network information) with an interaction term. In the cell line dataset, all metrics of each common nodes showed non-significant interaction terms suggesting parallel trends (Supplementary Data). We can observe visually that, for example, the interactions of GNL3 (9606.ENSP00000395772) do not change with time at all in any types of networks, while the interactions of MRPL9 (9606. ENSP00000357823) are not the same in the proteomic-based networks comparing the first and last generations, but the measured characteristics of the node overall remain the same (Fig. 5c). In the embryogenesis dataset, 28% of the common nodes’ degree centrality, 73% of the nodes’ clustering coefficient and 77% of the nodes’ betweenness centrality show non-significant slope suggesting parallel time-related changes of two of the three measured node characteristics (Fig. 5d, Supplementary Data). An example of a protein showing parallel time-related change in its clustering coefficient is Nt6b (7227.FBpp0309367), its clustering coefficient increases with time in both the transcriptomic-based and proteomic-based networks, but for example, the Fkbp59 (7227. FBpp0309663) protein shows opposite time-related trend in the two types of networks (Fig. 5e). It can also be seen in the examples that most changes already happen in the first three hours of development as we observed previously too.

Overall, the value of the rescaled node-level metrics suggests high similarity between the characteristics of the nodes in the transcriptomic-based and proteomics-based networks, but subtle differences may be present. The comparison of the time-related dynamics showed that while in the cell line dataset the time-related trend as well as the rate of change of the node properties are mostly similar, in the embryogenesis dataset, the time-related dynamics of the nodes’ degree is typically different, but the dynamics of the nodes’ clustering coefficient and betweenness centrality are mostly similar.

## 4 Discussion

We presented a comparative analysis of sample-specific protein-protein interaction networks estimated from transcriptomic and proteomic measurements. Although gene expression is widely considered a weak predictor of protein abundance, we found that both types of data provide similar insights into the complex system of protein-protein interactions in individual samples. While the coverage difference between the transcriptomic and proteomic data results in interaction networks at different resolutions, after correcting for the size, the characteristics of the two types of networks are barely distinguishable. We also found that the change in the network properties with time typically exhibits the same trend, though the extent of similarity was dataset-dependent. The difference between transcriptomic and proteomic data was revealed by the analysis of the rate of changes in characteristics: proteomic data-based networks showed larger deviation over time, while transcriptomic data-based networks exhibited less notable changes. We further found that the node characteristics are also similar, but some properties, such as the degree, tend to change differently with time in the two types of networks, while others, like the betweenness centrality, show larger similarity in the rate of changes too. These results suggest that while transcriptomic and proteomic data provide similar insights into the sample-specific PPIs, limitations of each omics should be kept in mind when interpreting the results, especially in the case of longitudinal trajectories.

Local, node-level characteristics provide further, more detailed insight into the network structures, revealing non-homogeneous time-related changes in the networks, which might bias the global characteristics. For example, the network-level average node betweenness centrality showed a different trend in the embryogenesis dataset, but the local structure revealed that there is still a high level of similarity among the betweenness of nodes. This points to a limitation of global network metrics and draws attention to the need for local characterization as well. We observed that the conclusions based on the two datasets analyzed in the present study only partially agree, which urges further, large-scale investigations involving additional datasets. It is important to explore whether the similarity of the two types of PPI network approximation is universal or depends on the context. For example, our findings in the cell line data completely support the findings of the original study presenting the data (Lu et al. 2023), namely that both omics were stable during generations in the cell line, and the network metrics also seem stable. On the other hand, it is believed that the transcriptome and the proteome significantly decouple during aging (Wei et al. 2015; Kelmer Sacramento et al. 2020), and it may increase the dissimilarity between the inferred interaction networks as well. However, we found that in the Drosophila embryogenesis dataset (Becker et al. 2018) the overlap of the networks inferred from transcriptomic and proteomic data increases with time, and many metrics also become more similar after the first 2-3 hours, which may suggest the opposite of decoupling during early development. On the other hand, previously, limited correlation was observed between the transcriptomic and proteomic measurements of this dataset (Becker et al. 2018), and the inferred networks also show divergence to some extent, but they still have a high level of similarity. An interesting open question is whether the larger deviation observed among the proteomic data-based networks is due to the relatively low coverage making small changes more visible, or it is due to the less stable nature of the proteome.

We observed a spectacular change (monotonic increase or decrease) for most of the network metrics during the first 3 hours of Drosophila embryogenesis, similarly to the original findings about the transcriptome and proteome (Casas-Vila et al. 2017; Becker et al. 2018), followed by no remarkable change in the remaining stages. The 3^rd^ hour, where this clear shift is observed, is the stage of gastrulation, where we previously revealed the rejuvenation of mouse and human embryos at the epigenetic and transcriptomic levels (Kerepesi et al. 2021; Trapp et al. 2021; Kerepesi and Gladyshev 2023; Zakar-Polyák et al. 2024). Accordingly, we hypothesize that the fruit fly embryo also rejuvenates during the stage of gastrulation; however, more investigation is needed to confirm it.

An important limitation of the present work is the static PPI network underlying both the transcriptomic-based and the proteomic-based approximations. The choice of the confidence threshold to include an interaction largely affects the prior network structure, and choosing a high confidence level might also exclude recently found interactions. Hence, it is important to further explore the effect of the threshold on the network structure and if also affects the similarity between the different approximations.

Overall, this work hopefully draws attention to the potential lying in the study of sample-specific interaction networks, which is still an underrepresented area in network biology. It also aspires to drive future works to discuss and highlight the limitations of approximations such as the present PPI network estimations and encourage deeper analysis to address the discrepancy among different approximation approaches.

### Data and code availability

The cell line dataset of Lu et al. is available at GEO under accession number GSE234201 and at iProX under subproject ID IPX0006212001. The embryogenesis dataset of Becker et al. and Casas-Vila et al. is available at GEO under accession number GSE121160 and in PRIDE under project ID PXD005713.

The networks generated in this study and their calculated characteristics are available on Zenodo at <u>10.5281/zenodo.21129948</u>. The codes generated in this study are available on Github at https://github.com/polyake/sample_specific_ppi.

---

**Supplementary Figure 1.**
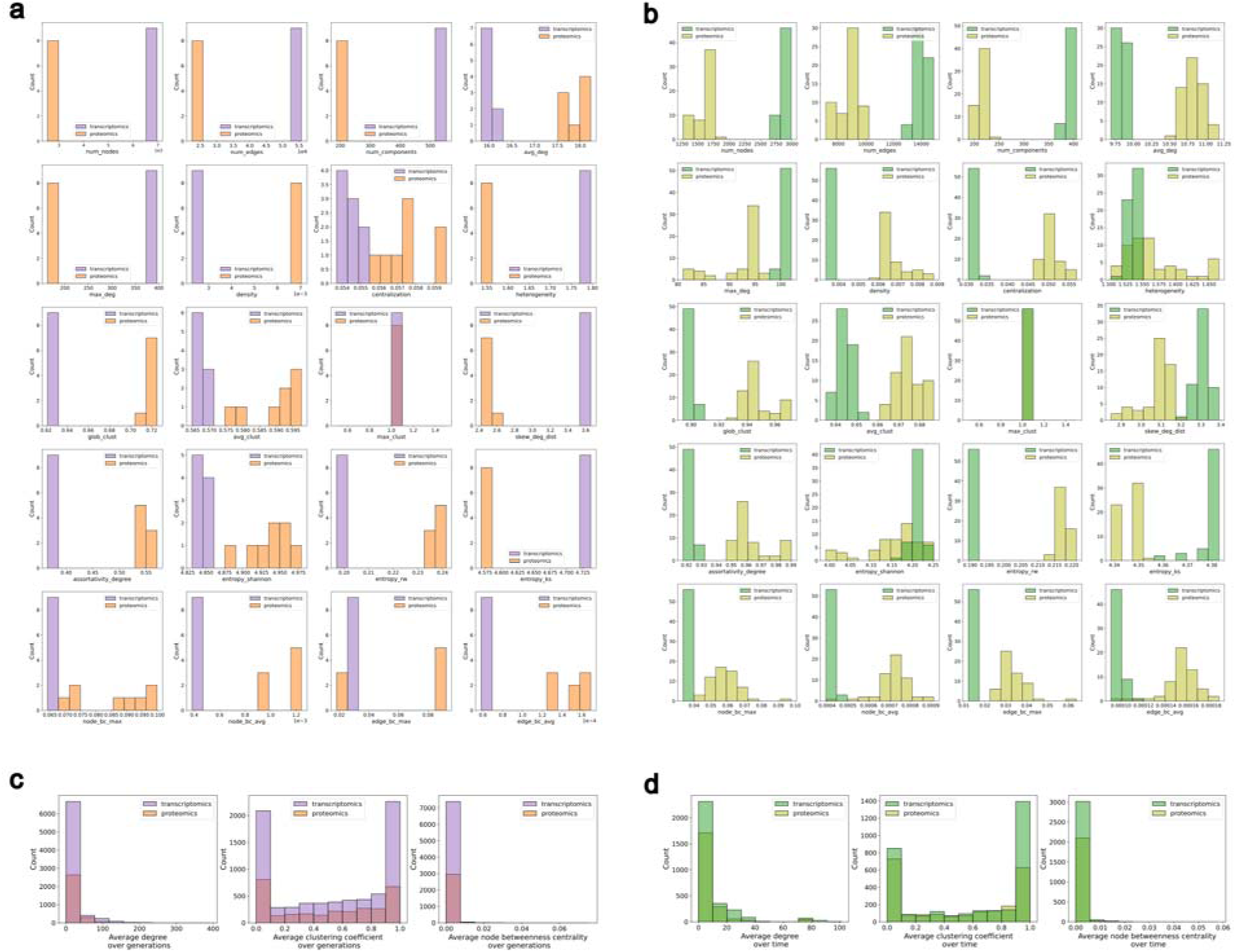
Original, non-scaled metric values of the networks inferred from transcriptomic and proteomic data. (a) Distribution of the non-scaled network metric values of the transcriptomic-based and proteomic-based networks in the cell line dataset. (b) Distribution of the non-scaled network metric values of the transcriptomic-based and proteomic-based networks in the embryogenesis dataset. (c) Distribution of the non-scaled node-level average metric values of the transcriptomic-based and proteomic-based networks in the cell line dataset. Average was calculated over generations for each node. (d) Distribution of the non-scaled node-level average metric values of the transcriptomic-based and proteomic-based networks in the embryogenesis dataset. Average was calculated over time points for each node.

**Supplementary Table 1.**
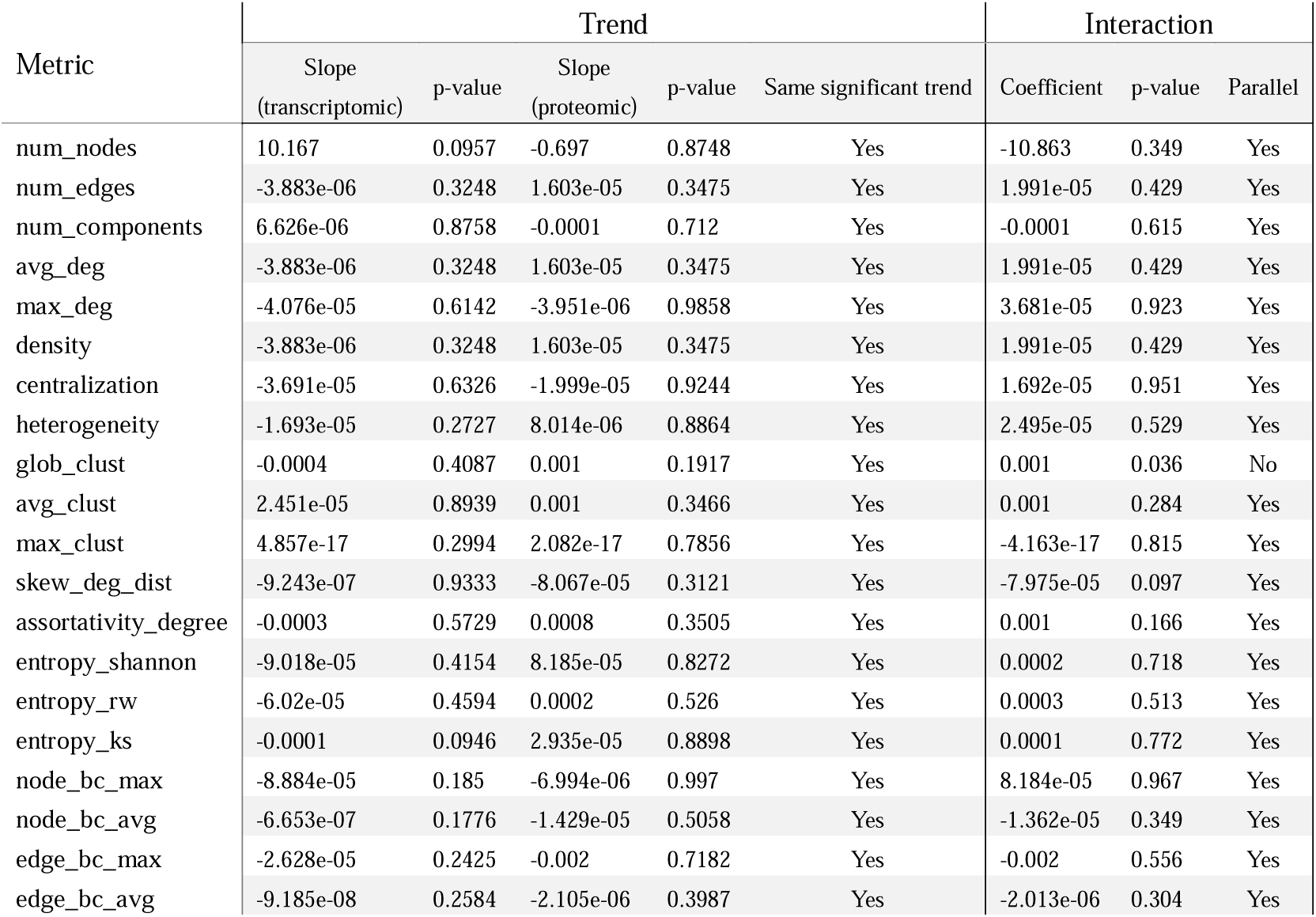
Summary of the time-related analysis results in the cell line dataset.

**Supplementary Table 2.** Summary of the time-related analysis results in the embryogenesis dataset.

| Metric | Trend |  |  |  |  | Interaction |  |  |
| --- | --- | --- | --- | --- | --- | --- | --- | --- |
|  | Slope<br>(transcriptomic) | p-value<br>(Bonf.) | Slope<br>(proteomic) | p-value<br>(Bonf.) | Same significant trend | Coefficient | p-value<br>(Bonf.) | Parallel |
| num_nodes | 5.388 | 2.8e-06 | 16.081 | 5.0e-10 | Yes | 10.693 | 0 | No |
| num_edges | -7.137e-06 | 4.0e-07 | -7.372e-05 | 3.0e-07 | Yes | -6.658e-05 | 0 | No |
| num_components | 6.846e-05 | 0.0227 | -0.0006 | 2.2e-07 | No | -0.0007 | 0 | No |
| avg_deg | -7.137e-06 | 4.0e-07 | -7.372e-05 | 3.0e-07 | Yes | -6.658e-05 | 0 | No |
| max_deg | -2.818e-05 | 0.0855 | -0.0003 | 7.4e-08 | No | -0.0005 | 0 | No |
| density | -7.137e-06 | 4.0e-07 | -7.372e-05 | 3.0e-07 | Yes | -6.658e-05 | 0 | No |
| centralization | -2.110e-05 | 0.3222 | -0.0002 | 1.6e-07 | No | -0.0002 | 0 | No |
| heterogeneity | -2.923e-05 | 0.0141 | -0.0005 | 2.4e-09 | Yes | -0.0004 | 0 | No |
| glob_clust | -0.0003 | 1.5e-07 | -0.0013 | 2.8e-11 | Yes | -0.001 | 0 | No |
| avg_clust | 0.0002 | 0.0708 | -0.0002 | 1 | Yes | -0.0004 | 0.66 | Yes |
| max_clust | -7.806e-18 | 1 | -1.084e-17 | 1 | Yes | -8.999e-17 | 1 | Yes |
| skew_deg_dist | -7.305e-06 | 1 | -0.0001 | 5.9e-14 | No | -0.0001 | 0 | No |
| assortativity_degree | -0.0001 | 2.1e-07 | -0.0007 | 2.9e-11 | Yes | -0.0005 | 0 | No |
| entropy_shannon | -8.627e-05 | 0.0026 | 0.0002 | 4.2e-05 | No | 0.0003 | 0 | No |
| entropy_rw | -5.988e-05 | 0.0022 | -2.252e-05 | 1 | No | 3.736e-05 | 1 | Yes |
| entropy_ks | -0.0001 | 0.0005 | -0.001 | 5.5e-08 | Yes | -0.001 | 0 | No |
| node_bc_max | -5.468e-05 | 0.0389 | -3.841e-05 | 1 | No | 1.627e-05 | 1 | Yes |
| node_bc_avg | -1.564e-06 | 2.6e-07 | 4.548e-06 | 0.0802 | No | 6.112e-06 | 0.04 | No |
| edge_bc_max | -3.041e-05 | 0.4253 | -0.0001 | 1 | Yes | -9.487e-05 | 1 | Yes |
| edge_bc_avg | -3.571e-07 | 1.5e-06 | 9.636e-07 | 0.0707 | No | 1.321e-06 | 0.02 | No |

